# Let Me Make You Happy, And I’ll Tell You How you look around: Using an Approach-Avoidance Task as an Embodied Emotion Prime in a Free-Viewing Task

**DOI:** 10.1101/2020.09.09.289249

**Authors:** Artur Czeszumski, Friederike Albers, Sven Walter, Peter König

## Abstract

The embodied approach of human cognition suggests that concepts are deeply dependent upon and constrained by an agent’s physical body’s characteristics, such as performed body movements. In this study, we attempted to broaden previous research on emotional priming, investigating the interaction of emotions and visual exploration. We used the joystick-based approach-avoidance task to influence the emotional states of participants, and subsequently, we presented pictures of news web pages on a computer screen and measured participant’s eye movements. As a result, the number of fixations on images increased, the total dwell time increased, and the average saccade length from outside of the images towards the images decreased after the bodily congruent priming phase. The combination of these effects suggests increased attention to web pages’ image content after the participants performed bodily congruent actions in the priming phase. Thus, congruent bodily interaction with images in the priming phase fosters visual interaction in the subsequent exploration phase.

## Introduction

Gaze-dependent shifts play a pivotal role in visual processing. Using modern eye-tracking techniques, it is possible to measure overt shifts of attention reliably and unobtrusively, helping us understand eye movement behavior. What one observes is influenced by at least three factors. First, attention is influenced by the external stimuli’ properties, processed in a bottom-up hierarchy (1, 2). This includes low-level features of the visual stimulus, for instance, contrast, contours, color, texture, and motion. However, it may also include more complex features like complex shapes of objects or the emotional valence of images (3, 4). Second, attention is influenced by internal variables like task-demands (5–7), as well as the observer’s emotional state (8). Third, the spatial factors like the central bias (9) and saccadic momentum (10) influence the selection of fixation targets. These three factors’ relative contribution is a matter of debate (11), and presumably depends on the precise circumstances (5). Additionally, to all different mentioned levels that attention can be influenced, it is crucial to operationalize attention itself (12). Our study used direction and allocation of eye movements to refer to attention (13).

When it comes to the role of emotional states affecting attention, it is useful to distinguish between an internal affective influence, e.g., the emotional state of the observer, and an external affective influence, e.g., the stimulus valence (8, 14–17). A situation in which attention is subject to both external and internal affective influences is when one explores web pages of online news portals. On the one hand, such web pages commonly contain positive alongside negative information, whereas, on the other hand, one is in a specific emotional state: Positive, negative, or neutral. Kaspar et al. (15) used such an environment to investigate the internal and external affective influences in a free-viewing task performed by young adults. The participants’ emotional state was primed by a series of either positively or negatively valenced visual stimuli. Subsequently, they had to explore web pages containing both positively and negatively valenced content. An analysis of the eye-tracking data revealed that a negative emotional state marginally elicited a more spatially extensive exploration and that attention for negative news increased in participants who were in a positive emotional state. Thus, the state of the observer and the external affective influence impacted the visual exploration.

The valence of the stimulus influences responses beyond visual exploration. Specifically, approach-behavior is naturally associated with an appraisal of something as ‘good.’ In contrast, avoidance behavior is naturally associated with an appraisal of something as ‘bad.’ As a consequence, we are faster (18) and more accurate (19) when making movements that correspond to their embodied meaning, i.e., approach for good, and avoid for bad. In particular, there is a general bodily tendency to approach positive and avoid negative cues and do so faster than vice versa (20). Moreover, it was shown that positive concepts and percepts are placed close-to-the-body locations, while negative concepts and percepts are placed away-from-the-body locations (21, 22). This gives evidence for a general bodily reaction to positive or negative stimuli (20, 23). For example, spider phobics avoid pictures of spiders more strongly than neutral cues (24); socially anxious people avoid smiling and angry faces faster than controls (25); schizophrenic patients with higher levels of oxytocin avoid angry faces faster than controls (26); individuals with anorexia-nervosa exhibit a decreased approach bias for food cues (27); and healthy adults pull positive words faster towards them while pushing negative words faster away (18). Thus, the valence of stimuli has a widespread impact on bodily states and actions.

The cognitive mechanisms of the automatic approach bias are still debated. One approach, the concept of embodied cognition, rejects the idea that an agent’s cognitive life can be understood without considering the particular morphological, biological, and physiological characteristics of its body (28–30). For instance, language processing (31), memory (19), visual-motor recalibration (32), or distance estimation (33) all rely on specific body characteristics. Moreover, even our abstract concepts are bodily ‘grounded’ and arise from the body (34). That is to say that according to the embodied approach of cognition and affectivity, cognitive and affective phenomena can be fully understood only by taking into account the specific morphological, biological, and physiological details of the agent’s body (28–30). In particular, bodily movements are specific to the kind of body we have, and to the environment, we interact with, and are thus naturally meaningful. Similarly, the embodied approach to cognition tries to explain the approach-avoidance behavior (35). Importantly for our study, affective states are also considered within the embodied cognition framework. (36) discuss emotions in relation to the body and beyond the body and the brain. Furthermore, (37) propose an action-oriented understanding of emotions.

However, although emotional priming has been an important topic in research on top-down influences in overt attention, in particular when it comes to disentangling external and internal affective influences, there is little research using embodied primes (38). As creatures with specific bodily morphology, our onto- and phylogeny make it natural that positive valence is pulling something towards us while pushing it away is negatively valenced (35). Since the human abdominal region is exceptionally vulnerable, we have to protect it by allowing only trustworthy objects to come close. Since survival requires energy, we have to pull nourishing objects towards us while avoiding rotten, poisonous, unsanitary, or noxious objects. While strangers must typically be kept at bay, procreation, nurturing infants, and giving them love and comfort require social approaching. The idea that approach- and avoidance-behavior is naturally associated with appraisals of something as ‘good’ or ‘bad’ is also in line with embodied accounts of emotions (36, 39), in particular with Damasio’s (40) ‘somatic marker’ theory, according to which emotions function to direct animals towards what is good and direct them away from what is bad. Hence, these considerations suggest that approaching something or pulling it towards us is naturally meaningful, indicating something is positive.

If the automatic approach bias is indeed a general bodily reaction to positive or negative stimuli (20), we should observe it in healthy adults performing an approach-avoidance task. This type of explanation raises new questions. Namely, if the bodily relation is crucial, we would expect an influence of the stimulus valence and a congruency effect. Body movements that are in line with our preferences (pull towards positive, “good”/push away negative, “bad”) should influence our eye movements differently than priming by incongruent actions preferences (pull towards negative, “bad”/push away positive, “good”). Therefore, we aim to answer that question with our design. The congruency effect would give support to the claim that embodied priming modulates our viewing behavior. That is, here, we are primarily interested in modulating the natural (embodied) action in response to a stimulus e.g., congruent vs. incongruent, as well as investigating the effects on subsequent visual exploratory behavior.

The present study builds on an embodied approach to the automatic approach bias in order to investigate (1) whether healthy adults exhibit a comparable automatic approach bias concerning positively and negatively valenced stimuli and (2) how a positive vs. negative emotional state, induced by a congruent vs. incongruent approach-avoidance task affects their overt attention in a free-viewing task.

## Methods

### Participants

Twenty participants (6 male, 18 right-handed, mean age of 22.6 years, standard deviation of 2 years) took part in the experiment. They gave written informed consent before the start of the experiment. Participants received either 9 C or course credits in exchange for their participation. All participants had normal or corrected to normal vision and were not aware of the study’s scientific purpose. They were either native German speakers or fluent in the German language. This was important since the presented stimuli included headlines written in German. The ethics committee of Osnabrück University approved the study.

### General Apparatus

We presented all stimuli on a 24” LCD monitor (BenQ XL2420T; BenQ, Taipeh, Taiwan) with a refresh rate of 114 Hz. Participants sat 80 cm away from the screen. The experiment was controlled by a PC (Dell) connected to an eye tracker computer via an Ethernet cable. We used a head-mounted eye tracker (Eye Link II system) from SR-Research Ltd. (SR-Research Ltd, Ontario, Canada) to track the participants’ eye movements. In turn, the eye tracker was connected to a DOS-based computer (Pentium 4; Dell, Round Rock, TX, USA) running the application software. In total, the eye tracker comprised three infrared cameras. The head camera recorded infrared sensors attached to the monitor’s corners to calculate the head position in relation to the screen continuously. This allowed a stable gaze recording irrespective of involuntary small head movements. The other two infrared cameras recorded the participants’ pupil positions. The sampling rate of binocular recordings was 500 Hz. The room was darkened during the entire experiment.

A 13-point calibration task preceded each recording. It consisted of fixation points appearing consecutively in random order at various screen locations, and participants were instructed to focus their gaze at these points. Each point had a visual angle size of 0.5°. We validated the calibration by calculating the drift error for each point. Thereby it was assured that the mean validation error stayed below a 0.3° visual angle and the maximum validation error below a 1° visual angle. The calibration was repeated until the mentioned accuracy was reached.

We used the eye tracker’s default settings to calculate saccades and fixations. Saccade detection was based on a velocity of at least 30° visual angle/s and acceleration of at least 8000° visual angles/s^2^. To trigger a saccade, the saccade signal had to be sustained for at least 4 ms. By the time the eyes moved significantly from the fixation point (i.e., exceeding a motion threshold), the saccade’s temporal and spatial onset had been defined. By default, we set this motion threshold to a 0.1° visual degree. After the saccade onset, the minimal saccade velocity was 25° visual degree/s. Following this, a period without a saccade was marked as fixation. Each trial was followed by a fixation cross appearing in the screen center to control drifts in measurements. The first fixation following each stimulus’s onset was excluded from our analysis because this was an artifact from the drift correction before the respective trial’s onset.

The joystick used for the approach-avoidance task (Logitech Attack TM 3; Logitech, Apples, Switzerland) was connected to the computer screen. Matlab’s Psychtoolbox V3 (r2017a; MathWorks Company) enabled us to record response times (pushing/pulling movements). The joystick was placed on the table in front of the participants. We used MATLAB to preprocess eye-tracking data and R to analyze all data. Analysis scripts and data are available online (https://osf.io/cyz9b/

### Stimuli

The experiment included two separate phases (see below for details). First, participants performed an approach-avoidance task, viewing isolated images. Afterward, participants visually explored web pages, each containing two embedded images with additional text columns in a typical newspaper layout. As we investigated the influence of the approach-avoidance task on later visual exploration, we labeled the isolated images as “primes”.

In this study, we used 88 full-colored images from the International Affective Picture Set (IAPS) (41). Kaspar et al. (15) used the identical stimulus set. Half of the images had a valence rated below 3 (IAPS scale) and served as negative primes. The other 44 images had valence ratings above 7 and served as positive primes. To prevent the images from blurring, we presented all of them in their native resolution of 1024 × 768 pixels on a gray background (RGB values: 182/182/182), centered in the middle of the screen (resolution of 1920 × 1080 pixels, Figure 2A).

In the present study, twenty-four prototypes of news web pages were used, previously designed by Kaspar et al. (15) (Figure 1). The web page images’ resolution fits the screens’ resolution (1920 × 1080). Two target areas, embedded by several textual and pictorial components, were constructed in one web page design (Figure 1A). Each main news article included either a negative or positive IAPS image (615 × 411 pixels), a matching heading, as well as a link to the entire news report. It is important to note that there was no other textual content regarding the main news. This was done to avoid attraction biases because of how appealing news may have been for individuals who participated. Since participants did not interact with the web pages, the link served no function, except for creating a realistic version of a news web page that can be found on the world wide web. The structure and content of the web pages remained the same throughout the whole study. However, the side of the negative and positive content was counterbalanced (Figure 1B, C). The 48 images, embedded in the 24 web pages, differed from those used in the priming sessions.

**Fig. 1.**
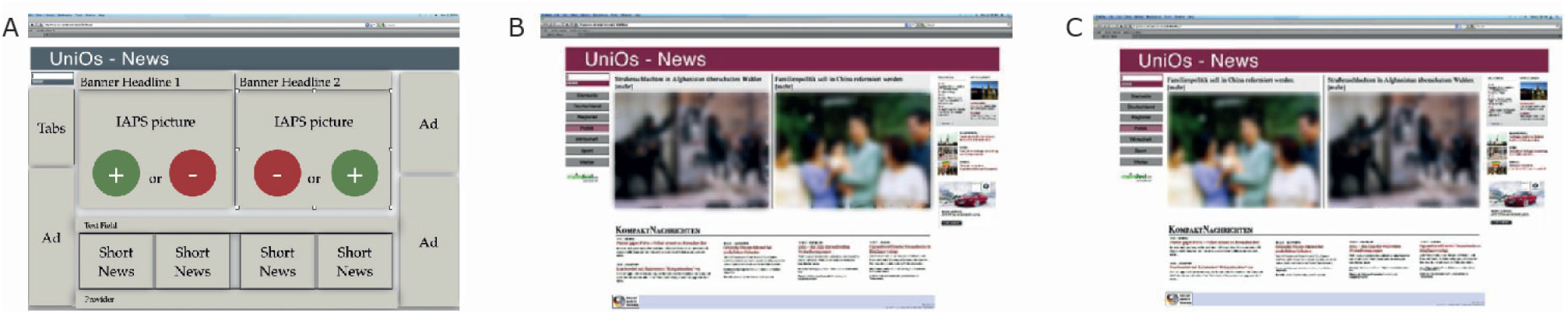
Web pages used in the study. (A) Prototype web page with different sections: headlines, adds, tabs, news, and images. (B) Example of a web page with a negative image on the left side. (C) Example of a web page with a negative image on the right side (reverse of (B)).

The additional elements on each web page were four short news reports about ongoing current affairs. These elements were placed below the main news articles. The frame around the main news articles was completed by flanking advertisements on the left and right sides (Figure 1A). Please note that the statistical properties of forward directed saccades and backward directed saccades (regressions) while reading the text do not enter the analysis presented here in any form. As a standard feature on regular web pages, the upper left corner was secured for a tabs region which is necessary for general navigation. Previous work by Kaspar et al. (15), using the same set of web pages, tested for the possibility that differences in eye movement parameters, within positive and negative images, could evolve from systematic differences in visual saliency. Therefore, a standard algorithm by Itti et al. (42) that extracts the physical features of images and, based on this, predicts fixation patterns was applied. In addition to this, a graph-based visual saliency (GBVS) developed by Harel et al. (43) was applied, as it predicts the fixations with a higher probability. After application, no difference regarding the visual saliency was found between the positive and negative images in the stimulus set (both *t*(35) ≤ 0.941, *p* ≥ 0.356).

### Procedure and Design

We divided the participants randomly into two groups. One group started with the congruent block of the approach-avoidance task. The other group started with the incongruent block. In each condition, participants faced a random sequence of 44 images of different valence (22 positive and 22 negative images). As soon as an image was presented, the participants had to respond with the joystick. The task paradigm required participants to push or pull the joystick in response to the image’s valence. Participants used their dominant hand to manipulate the joystick in front of them. Participants in the congruent task condition had to pull (approach) the joystick towards themselves whenever a positive image was shown, and push (avoid) the joystick whenever a negative image was shown. In the incongruent condition, participants had to act reversely. They had to pull the negative images towards themselves and push away the positive ones (Figure 2B, C). They were instructed to respond as quickly and as accurately as possible. It was not possible to rectify and correct response mistakes. Additionally, while moving the joystick towards or away, the image changed in size. The zoom feature of the approach-avoidance task was programmed in MATLAB’s Psychtoolbox V3 (r2017a; MathWorks Company); as such, a shown image smoothly decreased in size as soon as the joystick was pushed (Figure 2B). Conversely, the image size increased once the joystick was pulled (Figure 2C). It is important to note that participants were instructed to push or pull the joystick to its limit. Overall, participants took about 5 minutes to complete this first part of the experiment.

**Fig. 2.**
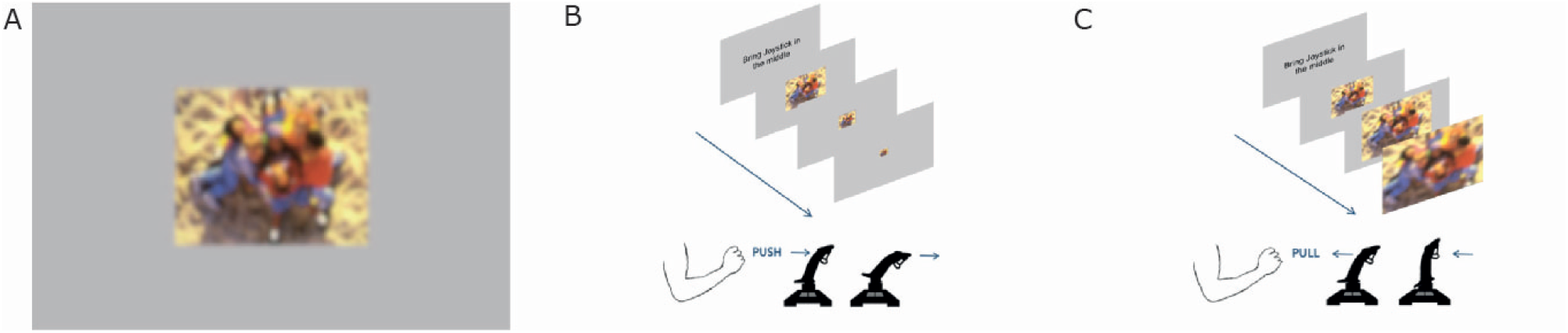
Embodied primes used in the study. (A) Example of one priming stimulus: IAPS image with surrounding gray frame. This photograph depicts positive content with children laughing and playing on a sandy playground. Hint: Due to restrictions of showing IAPS images in public, it is masked with a diffusing filter. (B) Incongruent condition: joystick pushed forward - positive prime zooms out. (C) Congruent condition: joystick pulled backward - positive prime zooms in.

In the subsequent eye-tracking session, we recorded the viewing behavior on prototypes of 12 news web pages. Following earlier research of Kaspar et al. (15) and ensuring the same experimental design, each web page was displayed for 15 seconds. We instructed participants to explore the web pages freely (free-viewing task).

After the eye-tracking session, participants had a short break. The second part of the experiment, directly after the break, required the participants to complete the joystick approach-avoidance task in the other condition. Participants who performed in the congruent approach-avoidance task had to complete the incongruent condition. The opposite applied to participants in the other group. After the second priming session, an additional 12 web pages were displayed following the same procedure described above (free-viewing task).

## Results

### Performance in the embodied approach-avoidance task

For the priming part of the experiment, we first calculated the accuracy of performance to check whether participants followed instructions. In the congruent condition, they had to pull positive and push negative primes. In the incongruent condition, the assigned actions were reversed. We found that participants made a low amount of errors (3.6%). This suggests that instructions were clear, and participants followed them. Therefore, we excluded error trials from any further analysis.

Second, we focused on response times in the experiment’s priming part to check whether positive and negative images in congruent and incongruent conditions involve different cognitive processes and, therefore, longer/shorter response times. As the data showed a skewed distribution, we log-transformed it before further analysis (we used natural logarithm)(44). We used a linear mixed model (LMM) to analyze response times. The LMM was calculated with the lme4 package (45), and *p*-values were based on Wald’s T-test using the lmerTest package (46). Degrees of freedoms were calculated using the Satterthwaite approximation. We modeled response times by image valence (positive and negative), and experimental condition (congruent and incongruent movements) as fixed effects and interactions between them. As random effects, we used random intercepts for grouping variable participants. For all predictors, we used effect coding scheme with binary factors coded as –0.5 and 0.5. Thus, the resulting estimates can be directly interpreted as the main effects. This coding scheme’s advantage is that the fixed effect intercept is estimated as the grand average across all conditions and not a baseline condition average. We found the main effect of the image valence (*t*(1673.02) = −3.726, *p* < .001) on response times (Figure 3B). The natural logarithm of the response time to negative stimuli was about 0.049 times smaller than to the positive stimuli. This corresponds to a speedup (reduction of response time) by a factor of 5.03%. Furthermore, we found the main effect of the congruency of the task (*t*(1673.04) = −4.71, *p* < .001) (Figure 3A). The response in the incongruent condition was slower by about 6.39%. The interaction between these effects was not significant (*t*(1673.02) = −0.87, *p* > .38). These results demonstrate independent additive effects of faster movements under congruent conditions and faster movements in response to negative pictures.

**Fig. 3.**
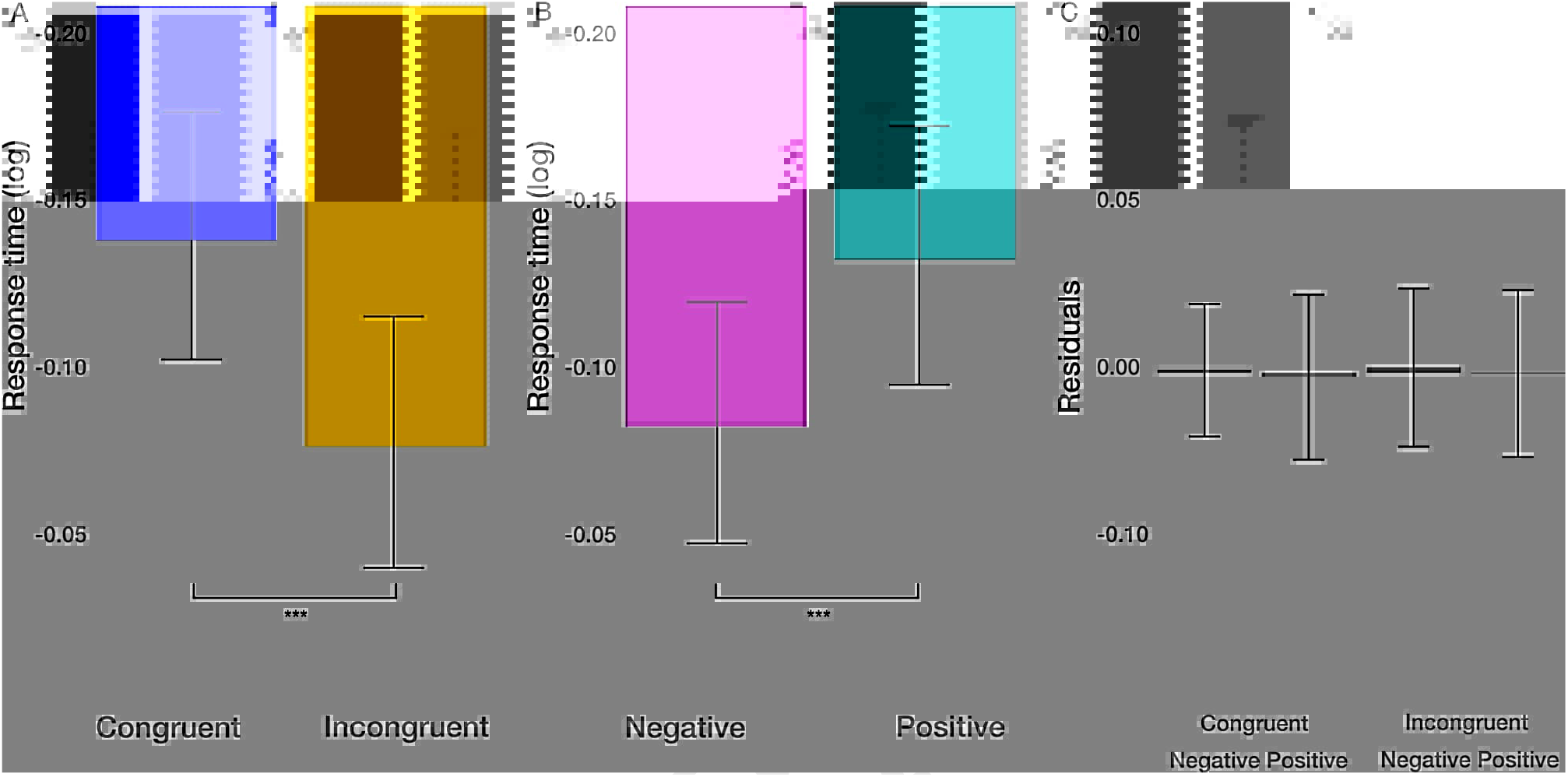
Response time results. (A) The main effect of the congruency of the priming. (B) The main effect of image valence (negative vs. positive). (C) Averaged residuals of all condition combinations from response time linear mixed model. In all plots, bars represent mean values and whiskers standard error of the mean.

### Eye movements in the free-viewing task (web pages)

As a next step, we investigated the effect of priming (condition: congruent and incongruent) on the viewing of news pages containing emotional stimuli (valence: negative and positive) on either side (side: left and right). The participants freely viewed different web pages containing one positive image and one negative image and additional filler texts, while we collected eye movement data. We characterized the exploration of these web pages with the two images as regions of interest (ROIs) with a various eye movement measures. Specifically, we used four different measures to quantify eye movements within ROIs: the average fixation duration within each image, the number of fixations within each image, the total dwell time on each image, and the length of saccades within each image. Additionally, we analyzed the number of saccades and their length from the outside to the inside of the images. For all six measures, we used the same statistical procedures. Similarly to the response time analysis, we employed linear mixed models. We modeled each of the variables by experimental condition (congruent and incongruent movements before the free-viewing task), image valence (positive and negative), and side of the image (left and right) as fixed effects and the interactions between them as random effects. We used random intercepts for grouping variable participants. For all predictors, we used effect coding scheme with binary factors coded as –0.5 and 0.5. We visually inspected the normality of the data. All variables, aside from dwell time, were log-transformed to achieve normally distributed data. Jointly, these measures and analyses allow the characterization of viewing behavior on the web pages after priming.

#### Fixation duration within ROIs

As the first measure, we used the average fixation duration within each ROI to measure the depth of processing (47). We did not find the effect of condition (*t*(8291.27) = −0.615, *p* = .54) on fixation duration. However, we found the main effect of the valence (*t*(8286.74) = −3.513, *p* < .001)(Figure 4A). The average fixation duration on negative images was longer by about 2.8%. Furthermore, we observed the main effect of the side (*t*(8284.26) = −3.093, *p* < .01)(Figure 4B). Fixations on the image displayed on the left side were longer by about 2.45%. Further, we found the significant interaction between valence and side (*t*(8283.33) = −3.318, *p* < .001)(Figure 4C). The difference in fixation duration on positive and negative images was larger on the right side. The size of this interaction was of the same order of magnitude as the main effect of the side of the image. That is, the longest average fixation duration was observed for the combination of negative images on the right side of the displayed web page. All other two-way and three-way interactions were not significant (*p* > .18). These results show that the displayed web page parameters, i.e., valence and side, had a significant influence on the depth of processing at individual fixation locations. However, this did not apply to the priming condition.

**Fig. 4.**
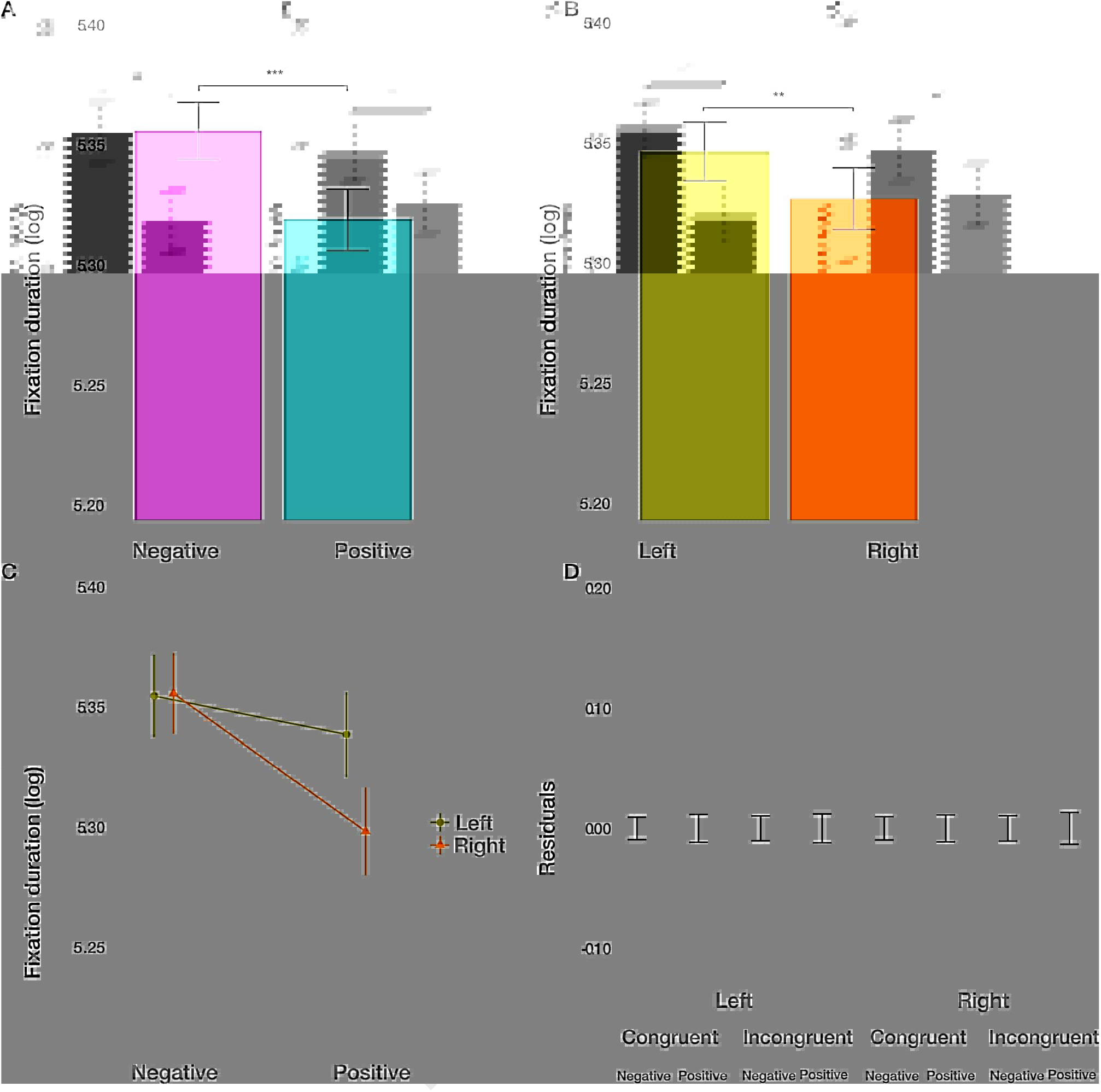
Fixation duration results. (A) The main effect of image valence (negative vs. positive). (B) The main effect of the side (left vs. right). (C) The interaction between valence and side. (D) Averaged residuals of all condition combinations from the fixation duration linear mixed model. In all plots, bars represent mean values and whiskers standard error of the mean.

#### Number of fixations within ROIs

Next, we considered the number of fixations within each image to measure attention devoted to the respective stimulus.

We found the main effect of condition (*t*(915.2) = −2.271, *p* = .0234)(Figure 5A). The number of fixations within the ROIs after congruent priming was larger by 11.47%. Furthermore, we observed the main effect of the valence (*t*(915.19) = −5.112, *p* < .001)(Figure 5B). Negative images captured 27.68% more fixations than positive images. Finally, we found the main effect of the side (*t*(915.17) = −2.816, *p* < .01)(Figure 5C). Images displayed on the left side captured 14.41% more fixations. All two-way and three-way interactions were not significant (*p* > .1). These results demonstrate additive effects, in terms of the logarithm of the number of fixations. Converted back to the number of fixations within the ROIs, this results in multiplicative effects on the condition, valence, and side on the attention devoted to the images.

**Fig. 5.**
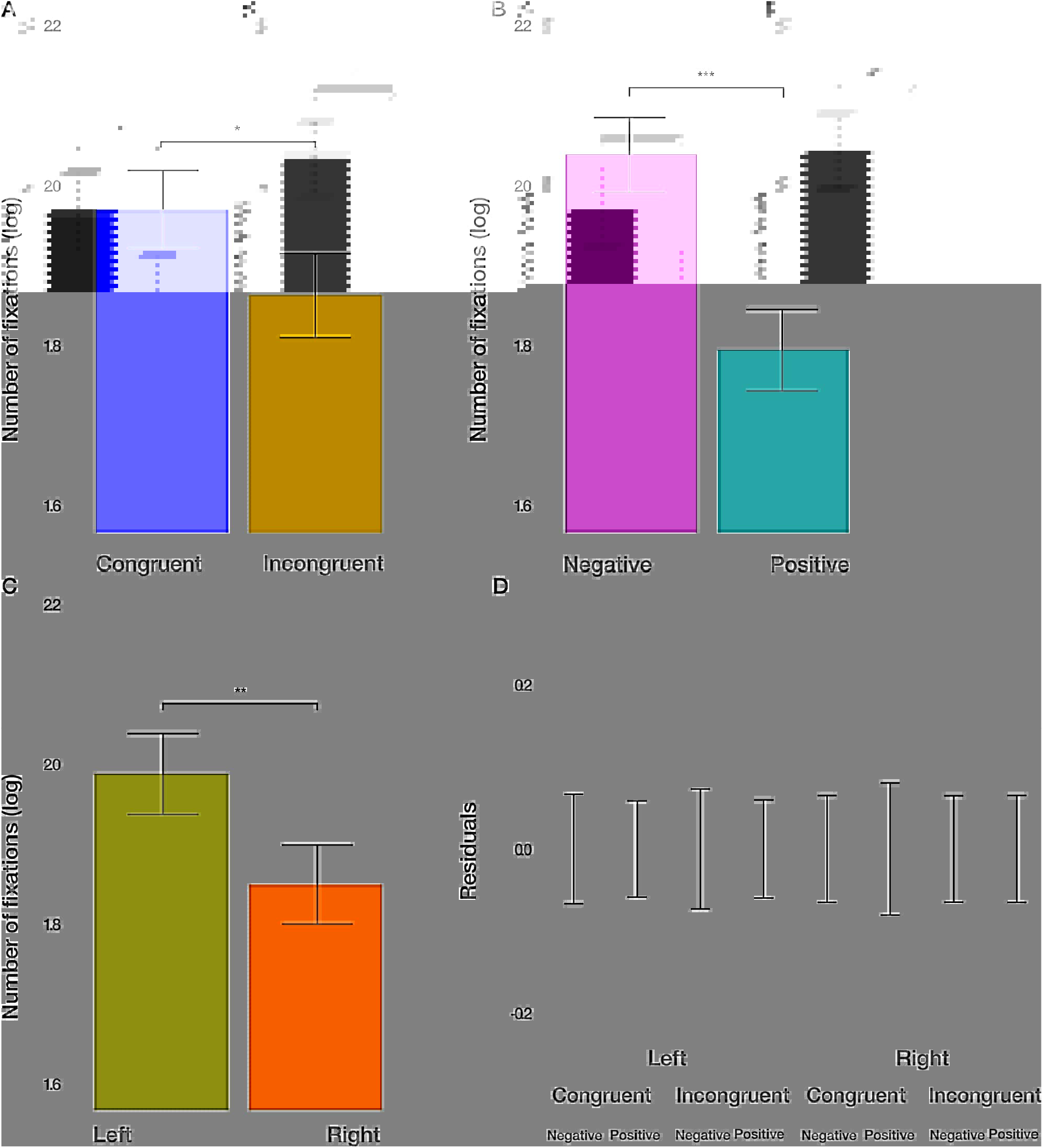
The number of fixations results. (A) The main effect of the congruency of the priming. (B) The main effect of valence (negative vs. positive). (C) The main effect of the side (left vs. right). (D) Averaged residuals of all condition combinations from the number of fixations linear mixed model. In all plots, bars represent mean values and whiskers standard error of the mean.

#### Dwell time within ROIs

The dwell time combines the aspects of fixation duration and the number of fixations within the ROIs. We found the main effect of condition (*t*(915.2) = −2.274, *p* = .0232)(Figure 6A). The dwell time within the ROIs after congruent priming was on average 190 ms larger. Furthermore, we observed the main effect of the valence (*t*(915.2) = −6.146, *p* < .001)(Figure 6B). Dwell time on negative images was on average 513 ms larger than on positive images. Finally, we observed the main effect of the side (*t*(915.17) = −3.381, *p* < .01)(Figure 6C). On average, the dwell time within images was on aver-age 282 ms larger on the left side. All two-way and three-way interactions were not significant (*p* > .33). These results resemble the results in the analysis of the number of fixations within ROIs. They provide evidence for independent effects of the priming condition, the valence of the viewed image, and the side of image location on the dwell time.

**Fig. 6.**
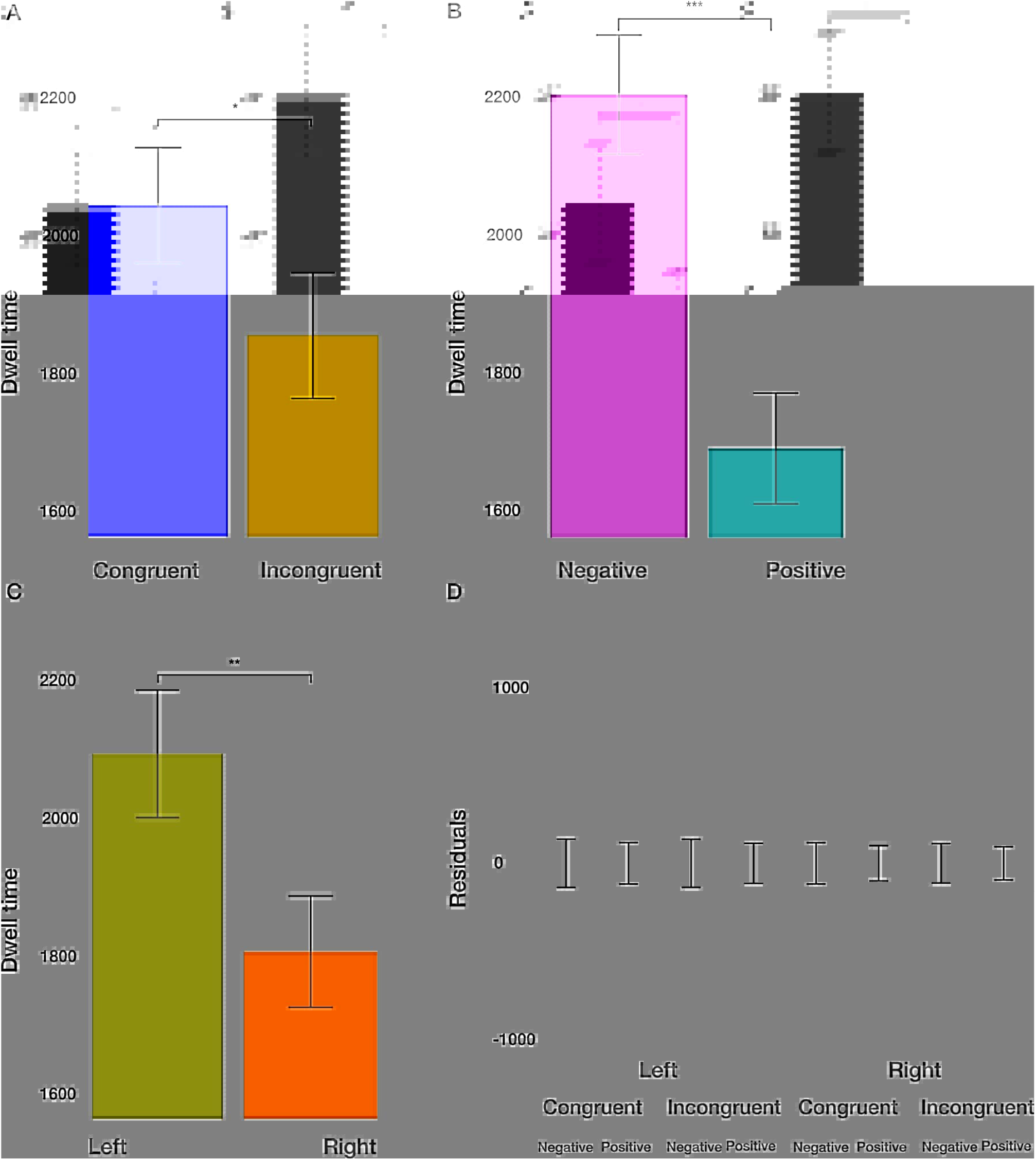
Dwell time results. (A) The main effect of the congruency of the priming. (B) The main effect of image valence (negative vs. positive). (C) The main effect of the side (left vs. right). (D) Averaged residuals of all condition combinations from the dwell time linear mixed model. In all plots, bars represent mean values and whiskers standard error of the mean.

#### Saccade’s length within ROIs

As a measure of exploration within the images, we used the saccadic length. We did not find the main effect of condition on the saccadic length (*t*(8296.9) = 1.321, *p* = .187). However, we did find the main effect of the image valence (*t*(8291.31) = 4.253, *p* < .001)(Figure 7A). Within negative images, saccades were shorter by 9.55%. Further, we observed a small but significant main effect of the side (*t*(8287.02) = 2.618, *p* < .01)(Figure 7B) on the saccade’s length. Saccades were shorter by 5.76% on the left side. Furthermore, we found significant two-way interactions between the image valence and the side of the image (*t*(8285.35) = −2.038, *p* = .0415)(Figure 7C), with a slightly larger difference in the saccadic length for positive and negative images on the left side. Additionally, we observed the interaction of condition and side (*t*(8289.3) = 2.907, *p* < .01)(Figure 7D).

**Fig. 7.**
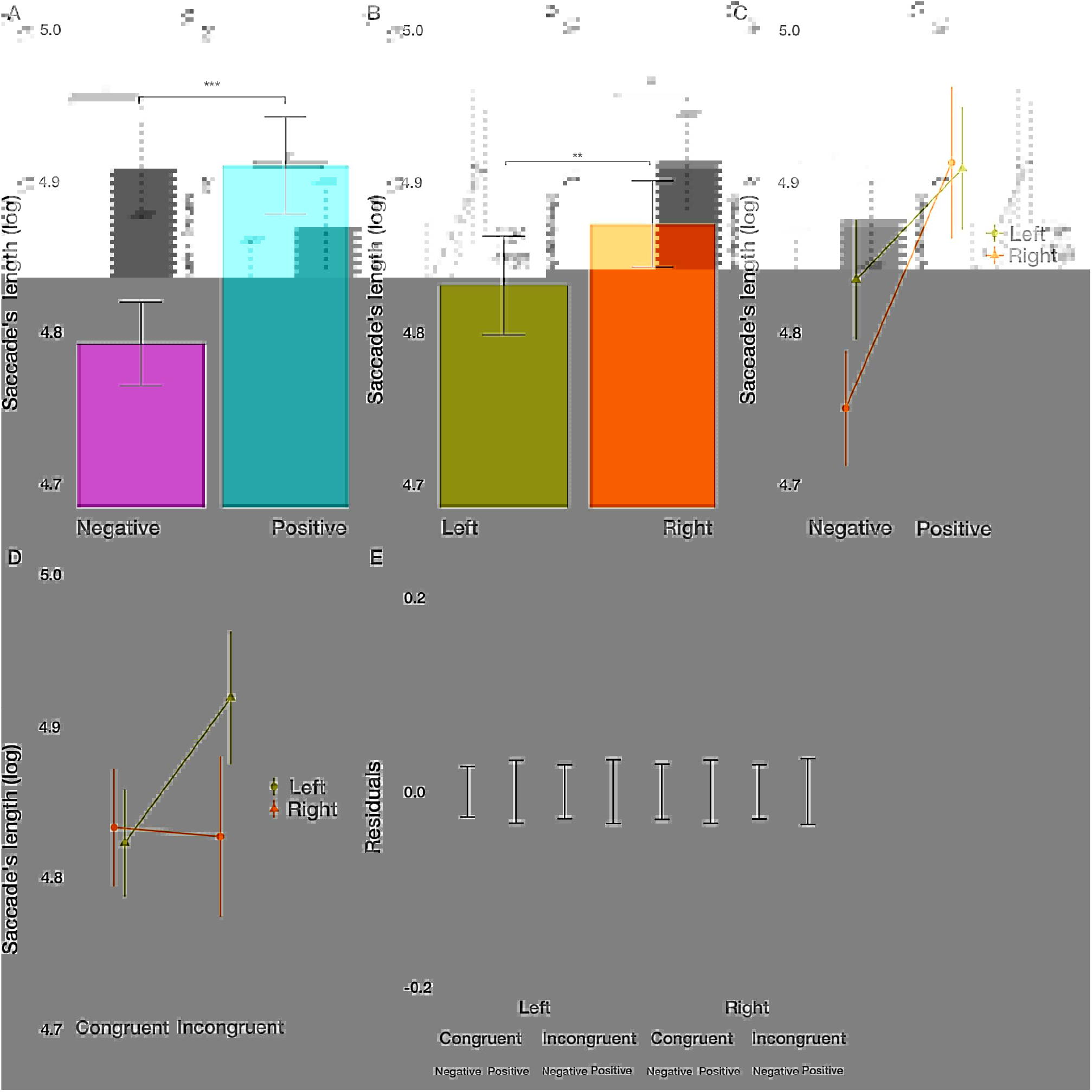
Saccade’s length results. (A) The main effect of image valence (negative vs. positive). (B) The main effect of the side (left vs. right). (C) The interaction between image valence and side of the image. (D) The interaction between congruency of the priming and side of the image. (E) Averaged residuals of all condition combinations from the saccade’s length linear mixed model. In all plots, bars represent mean values and whiskers standard error of the mean.

**Fig. 8.**
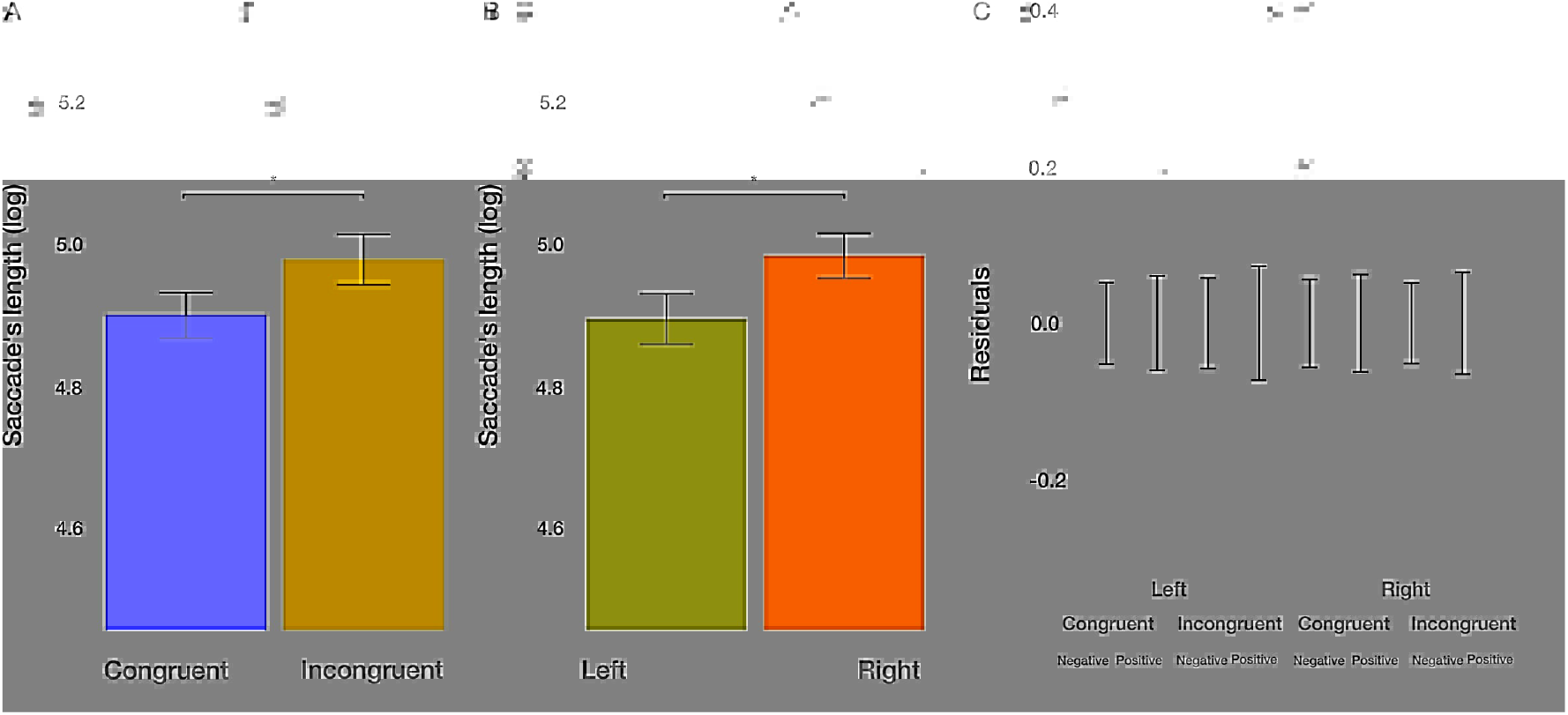
Saccade’s length from outside ROIs towards ROIs results. (A) The main effect of the congruency of the priming. (B) The main effect of the side of the image (left vs. right). (C) Averaged residuals of all condition combinations from the saccade’s length from outside ROIs towards ROIs linear mixed model. In all plots, bars represent mean values and whiskers standard error of the mean.

Whereas images displayed on the left condition were trivial, images on the right side were explored by longer saccades after incongruent priming. We did not find the interaction between condition and valance (*p* > .96), as well as no three-way interaction between all factors (*p* > .27). These results give evidence for a more focused exploration of images with negative valence, specifically when displayed on the left side. The priming condition modulated the influence of the side with a larger differential effect on the exploration of images displayed on the right side.

#### Number and length of saccades from outside ROIs towards ROIs

As a measure of how well the images can attract attention, we utilized the number of saccades from outside the image towards the inside. We found a trend towards significance for the main effect of the congruency of the priming (*t*(850.76) = −1.894, *p* = .0586). Participants, on average, made 6.96% more saccades from outside into an image after congruent condition. The main effect of the valence of the image (*t*(851.31) = −1.936, *p* = .0532) on the number of saccades from outside images towards them missed the significance threshold. Nominally, participants made 7.13% more saccades on negative images. The effect of the side of the image and all two-way and three-way interactions were not significant (all *p* > .12). Furthermore, we analyzed the length of saccades from outside images towards them. We found the main effect of congruency of the priming (*t*(2011.69) = 2.189, *p* = .0287). After priming with congruent actions, the average saccade length from outside into the images was shorter by 9.71%. Furthermore, the main effect of the side of the image (*t*(2009.88) = 2.269, *p* = .0234) on the saccade’s length from outside images towards them was significant. Saccades targeting the left image were, on average, shorter by 10.07%. The effect of the image valence and all two-way and three-way interactions were not significant (all *p* > .06). These results suggest that participants, on average, make shorter and more saccades after congruent priming from outside images towards them. Furthermore, on average, participants make longer saccades from outside images towards the images on the right side.

## Discussion

In the present study, we used an active bodily interaction with affective stimuli in an approach-avoidance task to investigate the influence on the later free visual exploration of news web pages containing emotional images. First, positively or negatively valenced images were zoomed in or out by pulling or pushing the joystick. Here we found multiplicative effects of valence and condition. That is, we could replicate the results of previous studies and report a faster response in the congruent condition. Furthermore, negative stimuli were reacted to faster. The lack of interaction and the additive effects on the log-response time suggest independent multiplicative effects on the base response time. Second, concerning the influence of embodied priming on eye movements in the subsequent free-viewing task, we observed the main effects of valence, side of the presentation, condition, as well as specific interactions. This demonstrates the influence of stimulus properties (valence), internal variables (priming by condition), and spatial properties (side) on visual exploration.

The following discussion will address the two main parts of our analyses. We will first discuss the viewing behavior (fixations and saccades) made only in the emotion-laden main news. The subsequent part deals with the approach and avoidance behavior while performing the approach-avoidance task while taking the response times into account. First, we observed an influence of the priming condition, i.e., performing congruent or incongruent actions in the approach-avoidance task on later visual exploration. Specifically, after priming in the congruent condition, the number of fixations on images increased, the total dwell time increased, and the average saccade length from outside of the images towards the images decreased. The combination of these effects suggests increased attention to web pages’ image content after the subjects performed congruent actions on images in the priming phase. Thus, congruent bodily interaction with images in the priming phase fosters visual interaction in the subsequent exploration phase.

Second, we found systematic effects of the valence of images in the exploration phase. Specifically, on images with negative valence, the average fixation duration was prolonged, more fixations were completed, and the total dwell time increased. Particularly, the average length of saccades within the images of negative valence decreased. This combination of effects speaks for increased scrutiny of negative valenced images.

Third, we observed a lateral asymmetry in the visual exploration phase. On average, participants displayed longer fixations, more fixations, and longer dwell time on the left side. The change of the length of saccades within the images was significant but quantitatively not relevant. In contrast, the saccades’ length from the outside of the images towards the inside was shorter on the left side. The combination of these effects largely resembles the effects observed with respect to the priming condition and suggests increased visual interaction with stimuli presented on the left side. These results further corroborate results suggesting spatial biases in eye movements (48). This suggests that biases towards the left side influence not only the number of fixations but the major properties of visual exploration as well.

Finally, with respect to interactions of condition, valence, and side, it is noteworthy that there were only a few. For the number of fixations, dwell time, and the average length of saccades from outside to inside, we did not observe any 2-way or 3-way interactions, and the residuals after discounting for the main effects were relatively small. Only for the average saccadic length within the images did we observe an interaction of valence*side and condition*side and for the average fixation duration an interaction of valence*side. It appears that with respect to the saccadic length within the images for the left side, the valence is more important, and the priming condition less important for images on the left side. The fixation duration is less affected by image valence on the left side. Overall, it is striking that the effects of the three independent variables are largely independent, and the interactions are limited to a few aspects.

The results of this study suggest that approach and avoidance reactions in humans have a direct influence on attention allocation and gaze behavior. We used the embodied cognition approach and, more specifically, the approach-avoidance task to explore its effect on eye movements. This study adds to the limited amount of eye-tracking research that has dealt with the interplay of top-down influences and bottom-up features. To induce a positive or a negative emotional state, Kaspar et al. (15) had participants watch either positive or negative sequences of 44 full-colored images from the International Affective Picture System (IAPS; (41)) with a valence rating below 3 for negative and a valence rating above 7 for positive primes. In the subsequent eye-tracking session, they presented 24 similarly structured webpages that included a positive and a negative IAPS image: one on the left, the other on the right. They found that a negative emotional state marginally elicited a more spatially extensive exploration. In our study, we used the same news web pages. However, in-stead of inducing emotional states by passively watching pictures, we used an approach-avoidance task as an embodied prime for positive and negative emotional states. In contrast, no specific emotional valence was primed, but rather the congruent or incongruent action, i.e. approach/avoidance of positive/negative valenced stimuli or the reverse assignment. There is ample evidence that our emotions affect our visual behavior. Regarding the direct effect of emotions on visual exploration, the broaden-and-build model of positive emotions (49) claims that positive emotions such as joy, interest, elation, or love, temporarily expand the focus of attention, therefore, increasing the thought-action repertoire by fostering interest in the environment and encouraging play and exploration (50). Accordingly, being in a happy emotional state versus being in a sad or neutral emotional state has been shown to increase participants’ breadth of attention (51). Similarly, Wadlinger et al. (52) found that the distribution of participants’ fixations on an image is broader in individuals induced into a positive emotional state, with more frequent saccades to neutral or positively valenced parts and with more fixations on positively valenced peripheral stimuli. Whereas, broadened attention is often associated with anxiety (53), which has led some to speculate that this might be an adaption to a negative emotional state (54), while a positive emotional state may reduce the motivation to scrutinize the environment because of an increased feeling of safety (55). Part of the explanation for these diverse findings may be that the emotional state induction procedures are also diverse, particularly concerning neutral emotional states. For instance, whereas some actively induce a positive emotional state by offering participants a bag of candies but simply do nothing in the neutral condition (52), others rely on a waiting room manipulation to actively induce a neutral emotional state (56). Others have also been known to use movies (57) or music (58). According to Kaspar et al. (15), this diversity of emotional state induction method questions the assumption that a neutral emotional state is always an adequate control condition. This may help explain why being in a negative emotional state had the same effect as being in a neutral emotional state according to some studies (51). Whereas other studies found similar effects of being in a positive and neutral emotional state (59). In light of this still unresolved issue, the present study followed Kaspar et al. (15) and solely contrasted positive with negative emotional states and focused on the effects of priming in congruent vs. incongruent actions in an approach-avoidance task.

When investigating embodied cognition, a high degree of ecological validity is necessary. We instructed participants to use a joystick to either approach or avoid positively or negatively valenced pictures displayed on a screen (60). To increase immersion, we implemented a visual ‘zooming-effect’ (24). When the joystick was pushed, the images were zoomed out, and when it was pulled, the images were zoomed in. This not only ensured a more realistic impression of movement towards or away from the images, but it also illustrated any ambiguity in the participants’ arm movements. The appraisal of a movement depends upon what is achieved (61, 62). Stretching out one’s arm often indicates a negatively valenced avoidance-behavior, i.e., when a harmful object is pushed away. Yet, it can also be an indispensable part of a positively valenced approach-behavior, for instance when one reaches for nourishing food or one’s infant. A joystick-based Approach avoidance task with a zooming-effect resolves this ambivalence. To further increase the immersion utilizing techniques of virtual reality offer themselves.

In addition to the effect of participants’ emotional states on their attention, we also explored the approach and avoidance behavior in the priming conditions. Since the IAPS pictures have exhibited an impact on the emotions of participants and therefore serve as a reliable priming method (15), we made use of them in our study to also modulate congruent vs. incongruent actions by the participants. Previous study designs have solely presented participants with a row of pictures within one category. However, our study design differs from previous work in that we let the participants visually and physically interact with the depicted pictures. For this reason, we joined pictures of two valence categories in one task, which had to be treated differently. Since we were working with IAPS images, it is worthy of mentioning that the highly negative images were also accompanied by a higher level of arousal, in contrast to highly positive images (41). This applies to the IAPS images within the priming block and the images embedded in the news web pages. As Kaspar et al. (15) note, negative emotions, such as anxiety, anger, and fear, also happen to be more arousing for the participants compared to positive emotions, such as pride or happiness. This applies as well to negative and positive emotional conditions. As mentioned in the introduction, along with the increment of arousal in negative emotions comes an increase in attention. This is caused by the initiation of survival-related actions related to behavioral and physical fight-or-flight responses (50). In turn, the specific arousal, which is immediately elicited by the mere presence of the valenced images, is interwoven with the arousal that is elicited by the interactive treatment of the images in the priming condition. Since the primes used in this study comprised of both valence categories, it is challenging to make any explicit distinction. However, the approach-avoidance task served as an authentic method to strive for and impact the internal approach and avoidance reactions in humans. Participants in the incongruent priming condition were significantly slower in treating the primes as instructed. Thus, they were slower to pull negative images towards themselves and push positive images away from themselves. The difficulty of the incongruent priming condition task was reflected in the participants’ response times. In general, the task instruction (to pull negative images towards oneself) essentially acts against the avoidance effect, which has been presented as an example of embodied cognition that emphasizes action-oriented behavior, i.e., actions related to survival. A direct comparison of both task conditions clearly revealed the avoidance effect. In the congruent condition, participants were significantly faster to avoid the negative stimuli, compared with avoiding the positive stimuli in the incongruent condition.

In controlled attentional shifts, older adults show a positivity bias and negativity avoidance (63). In contrast, no such bias is observable for automatic attentional shifts (64–66). The results are inconclusive for younger adults. Some studies find a preference for negative stimuli (3, 67), while others report a tendency to avoid negative stimuli (68). Some studies find emotional state-incongruent preferences (69, 70), while others report emotional state-congruent preferences (71–74). Presumably, this inconsistency is partly because studies only focused on external affective influences and disregarded the participants’ emotional state. However, considering the participants’ emotional states is crucial because one’s emotional state can determine one’s current goals. In fact, when emotional regulation is the primary goal of younger adults, they focus less on negative images and more on positive images (75). Moreover, students who learn to focus on positive stimuli subsequently show reduced attention for negative stimuli (52), indicating that attention is a powerful tool for emotional states regulation (76). In contrast, Das et al. (77) found that a positive emotional state can increase attention for negative information. However, the primary goal of young adults exploring news pages is arguably not emotional state regulation. They are rather in a ‘browsing mode’ in which they search for personally interesting information. In such a mode, features of the stimulus, such as its valence, are more likely to catch the observer’s attention (78). In contrast to these mixed results, the effects of congruent vs. incongruent conditions in the present study are relatively straightforward.

Many studies investigated an automatic approach bias in patients with substance abuse disorders. Individuals with a substance abuse disorder exhibit an automatic bias toward drug-related words (79) or pictures (80). In stimulus-response compatibility tasks, in which participants have to use a joystick to move cues either away or towards themselves, they approach rather than avoid drug-related cues and they approach them faster than they avoid them. In an implicit approach-avoidance task, in which participants push and pull cues according to formal features (like the format of a picture (81) or its vertical alignment (82)), heavy drinkers (83), patients with alcohol abuse disorder (81, 84), heroin addicts (85), smokers (86–88), and cannabis users (82) approach drug cues faster than healthy controls. In an explicit approach-avoidance task in which participants either push away drug cues while pulling neutral cues towards them or vice versa, individuals with alcohol abuse disorder approach drug cues faster than they avoid them (89). Thus, there is accumulating evidence for a general automatic approach/avoidance bias related to substance abuse.

In summary, we present how congruent embodied priming influences eye movements in a free-viewing task. Results presented in our study suggest that prior congruent movements, in line with our bodily reactions, can influence how we scrutinize images presented on the World Wide Web.

We found that movements in line with our bodily reactions (approach positive and avoid negative) influence how we observe images presented on the World Wide Web.

## Author Contributions

Conceived the study (PK, SW). Study design (FA, PK), data collection (FA), data analysis (AC, FA, PK). Initial draft of the manuscript (AC). Revisions and finalizing the manuscript (AC, FA, PK).

## Acknowledgments and Funding

We gratefully acknowledge the support by the DFG-funded Research Training Group Situated Cognition (GRK 2185/1), “Niedersächsischen Innovationsförderprogramms für Forschung und Entwicklung in Unternehmen (NBank)-EyeTrax, and the Deutsche Forschungsgemeinschaft (DFG) Open Access Publishing Fund of Osnabrück University. Moreover, we would like to thank Clayton Thompson for feedback on the manuscript.

## Data Availability Statement

The datasets for this study can be found at https://osf.io/cyz9b/.

